# Hearing sounds when the eyes move: A case study implicating the tensor tympani in eye movement-related peripheral auditory activity

**DOI:** 10.64898/2026.03.24.713974

**Authors:** Cynthia D. King, Tingan Zhu, Jennifer M. Groh

**Affiliations:** Department of Psychology & Neuroscience, Duke University, Durham NC; Department of Neurobiology, Duke University, Durham NC

## Abstract

Information about eye movements is necessary for linking auditory and visual information across space. Recent work has suggested that such signals are incorporated into processing at the level of the ear itself (Gruters, Murphy et al. 2018). Here we report confirmation that the eye movement signals that reach the ear can produce perceptual consequences, via a case report of an unusual participant with tensor tympani myoclonus who hears sounds when she moves her eyes. The sounds she hears could be recorded with a microphone in the ear in which she hears them (left), and occurred for large leftward eye movements to extreme orbital positions of the eyes. The sounds elicited by this participant’s eye movements were reminiscent of eye movement-related eardrum oscillations (EMREOs, (Gruters, Murphy et al. 2018, Bröhl and Kayser 2023, King, Lovich et al. 2023, Lovich, King et al. 2023, Lovich, King et al. 2023, Abbasi, King et al. 2025, Sotero Silva, Kayser et al. 2025, King and Groh 2026, Leon, Ramos et al. 2026, Sotero Silva, Brohl et al. 2026)), but were larger and longer lasting than classical EMREOs, helping to explain why they were audible to her. Overall, the observations from this patient help establish that (a) eye movement-related signals specifically reach the tensor tympani muscle and that (b) when there is an abnormality involving that muscle, such signals can lead to actual audible percepts. Given that the tensor tympani contributes to the regulation of sound transmission in the middle ear, these findings support that eye movement signals reaching the ear have functional consequences for auditory perception. The findings also expand the types of medical conditions that produce gaze-evoked tinnitus, to date most commonly observed in connection with acoustic neuromas.

## Introduction

With some exceptions (e.g. ventriloquism) (Kording, Beierholm et al. 2007, Rohe and Noppeney 2015, Hong, Badde et al. 2021), it is normally easy to tell if a sound is coming from the same place as a visual stimulus. However, under the hood, the brain works hard to create this experience -- via a dynamic mapping between visual space, which is anchored to the retina, and auditory space, which is based on differences in sound arrival time, loudness, and frequency content across the two ears. The mapping must be dynamic because the eyes are in frequent motion – saccading about three times per second, covering a range of roughly 80 degrees – changing the relationship between the retina and the ears with every gaze shift.

It is truly remarkable that we are not normally aware of these eye movements – perceptual integration of visual and auditory space is seamless, automatic, and continuous in time despite such movements of the eyes. Early work concerning the neural basis of this eye movement dependent dynamic mapping between visual and auditory space focused on the superior colliculus in monkeys, and showed that auditory response patterns varied with the position of the eyes in the orbits (Jay and Sparks 1984, Jay and Sparks 1987, Jay and Sparks 1987, Hartline, Vimal et al. 1995, Zella, Brugge et al. 2001, Populin, Tollin et al. 2004, Lee and Groh 2012, Caruso, Pages et al. 2021). Subsequent work extended these findings not only to other oculomotor and association areas (Russo and Bruce 1994, Stricanne, Andersen et al. 1996, Cohen and Andersen 2000, Mullette-Gillman, Cohen et al. 2005, Mullette-Gillman, Cohen et al. 2009, Caruso, Pages et al. 2019, Caruso, Pages et al. 2021), but also to areas within the auditory pathway, such as the inferior colliculus (Groh, Trause et al. 2001, Zwiers, Versnel et al. 2004, Porter, Metzger et al. 2006, Bulkin and Groh 2012, Bulkin and Groh 2012, Willett, Groh et al. 2019) and auditory cortex (Werner-Reiss, Kelly et al. 2003, Fu, Shah et al. 2004, Maier and Groh 2010). The ubiquitous nature of these results in a wide range of brain areas prompted experiments to probe where signals related to eye movements *first* appear in the auditory pathway and led to the discovery of a novel form of otoacoustic emission, the eye movement-related eardrum oscillation, or EMREO (Gruters, Murphy et al. 2018, Bröhl and Kayser 2023, King, Lovich et al. 2023, Lovich, King et al. 2023, Lovich, King et al. 2023, Abbasi, King et al. 2025, Sotero Silva, Kayser et al. 2025, King and Groh 2026, Leon, Ramos et al. 2026, Sotero Silva, Brohl et al. 2026). These faint sounds can be recorded with earbud microphones in the ear canal, and are thought to be universally present in all humans with normal auditory function (King, Lovich et al. 2023).

However, while the hypothesis that these signals are the first step of a computational process involved in dynamically connecting the visual and auditory scenes despite eye movements is plausible, the lack of a direct perceptual signature of eye movements in connection to hearing has left this connection uncertain – the very success of the underlying computational process producing this dynamic mapping makes it somewhat difficult to verify whether signals at any particular stage of the pathway are used for this process.

Here we bridge this gap, via a case report of a human subject with a dysfunction involving a component of the peripheral auditory system. This patient can hear sounds when she moves her eyes in one particular direction. Her relevant history involves myoclonus, or spasm, in the tensor tympani middle ear muscle. We were able to record the sound that she reported hearing with eye movements, and confirm that it was larger and longer than the EMREOs recorded in participants in normal auditory function. These findings directly bridge for the first time an aspect of the peripheral auditory system (the tensor tympani) where EMREOs are observed with a perceptual experience associated with eye movements – in this case, an anomalous one.

## Results

### 1. Characterizing the audible sounds

**S98** is a 67 y.o. woman who reached out to us upon learning of our research concerning how eye movements are involved in hearing. She reported that she could hear a sound when she made certain eye movements, and that sometimes it was loud enough for her husband to hear it when standing close to her. She shared that she has been diagnosed with tensor tympani myoclonus, or spasm of the tensor tympani muscle. Superior semicircular canal dehiscence (SSCD), a common cause for hearing sounds during eye movements, was not considered the cause of her complaint as this condition typically includes other symptoms that she did not exhibit (see Methods). We invited her to visit the lab to test and quantify what she was hearing. At that time, screenings revealed normal hearing and tympanometric measures bilaterally.

Qualitatively, S98 described the sounds she hears as a kind of fluttering sound that she perceives in her left ear when she looks to the far left. It was not clear from her verbal report if she only hears the sound while the eyes were moving vs. while the eyes were stably fixating but in a leftward position, or both. Clarifying those details was one of the goals of testing.

At the time of her visits, she reported only experiencing the sound for particularly large eye movements, outside the range of saccades that we normally use for our standard visually-guided saccade tasks (+/-18° from central fixation). Accordingly, we asked her to sit in the sound booth and voluntarily make the eye movements that were associated with her eye movement-related sound percepts. Saccading from a fixation point (+30°) to the right to a position (-32°) to the left and slightly up (+6°) reliably produced the sound percepts. Of note, she could not perceive the fluttering sound when she initiated saccades from central fixation (0°, 0°) to (-32°, +6°,) indicating that the length, direction and target location of the saccade were all necessary factors in generating her perception of the fluttering sound.

To obtain recordings of the sounds she heard, we placed an earbud microphone (ER-10b+) in the left ear and recorded while she performed the saccade task. To obtain corresponding perceptual reports, we placed another microphone on the desk in front of her, and asked her to tap this microphone when she successfully elicited a sound percept by moving her eyes.

Eye tracking while performing this large saccade task was tricky: the video eye tracker (EyeLink 1000)’s software could not fully capture these eye movements across this expanse of the oculomotor range. However, the video display of the eye on the EyeLink computer did show the eye movements. Accordingly, we recorded the video display with a different camera and aligned the recorded video with the microphone recordings to ascertain the approximate timing of the sound percepts with respect to saccade onset/offset and periods of fixation, as manually identified by reviewing the footage.

Figure 1A shows the microphone recordings while S98 made saccades; the sounds recorded in her left ear are shown in blue, while the red trace shows the perceptual-report microphone. The putative sounds perceived by the subject can be seen as the elevated bursts occurring just prior to the peaks in the red trace. We played an amplified version of the left ear recording back to her, and she stated that they were like the sounds that she perceives.

**Figure 1.**
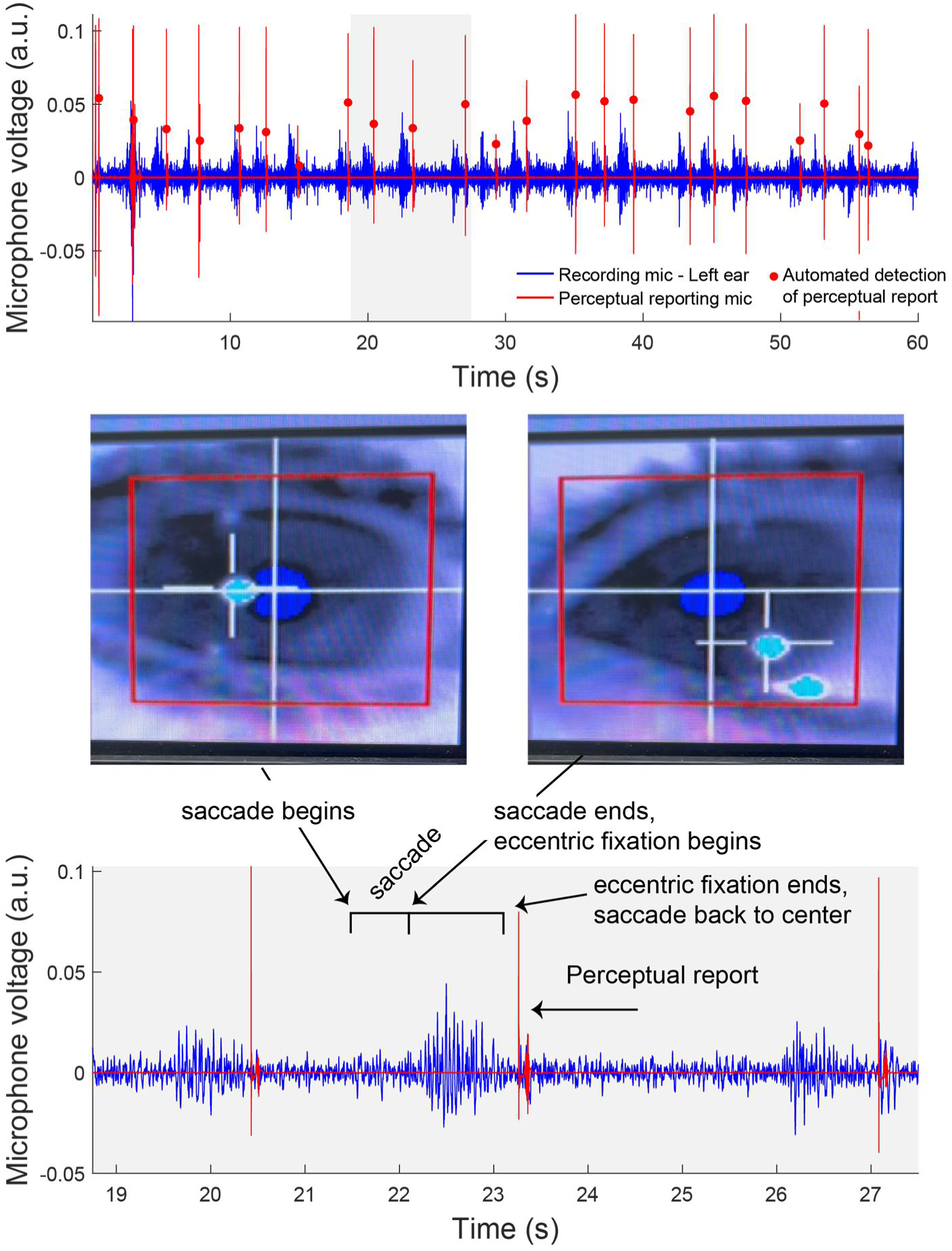
Recording of subject-perceived sounds related to extreme eye movements. A. Microphone recording during repeated large leftward eye movements, self-generated with the goal of eliciting the sound. The left-ear microphone recording is in blue. The subject tapped a second microphone to mark when she heard a sound; these tappings produced big spikes as shown in red. Automatic identification of the onset of each tap is marked with a red dot (“findpeaks” function, Matlab). B. The eye movements were saved by making a video of the eye as it was displayed on the screen of the EyeLink1000 computer. Audio and video recording were aligned to an estimated accuracy of 50 ms (limited by the monitor refresh rate of ∼70 Hz and video recording of 29 Hz; see Methods). Eye movement onset and offsets were then identified by visual inspection of this video-of-a-video. C. Enlarged view of a section of the recording (gray box from 1A) suggests that the audible sound was triggered by the saccade, lasted about 1 second, and occurred mainly while the eyes were holding in an eccentric fixation position.

It can be immediately inferred from the duration of these sounds (about 1 second) that they are not limited to occurring during the saccades only, as 62° saccades would normally take about 150-300 ms. Indeed, when the times of saccade onsets and offsets were marked by viewing the accompanying video (1B), it appears that the bulk of these particularly audible sounds occur during steady fixation at the far eccentric position (1C).

To more fully quantify the timing of the sounds with respect to saccades and fixation, we marked the beginning and end of the saccades from the rightward fixation to the leftward eccentric position as well as the onset of the saccade returning the eyes to a more central position, i.e. the end of eccentric fixation. Figure 2 shows the microphone traces associated with each reported sound across the recording aligned on the end of eccentric fixation. In some cases there was more than one saccade – if the subject didn’t elicit a sound with the first saccade, she would make a second one even farther to the left, and only return to a more central location after she had triggered her sound percept. When only one saccade occurred (Figure 2A), there was an evident sound beginning around the end of that saccade (orange vertical line) and lasting for about a second. When two saccades occurred (Figure 2B), the onset of the audible sound is more closely tied to the second saccade (orange dots).

**Figure 2.**
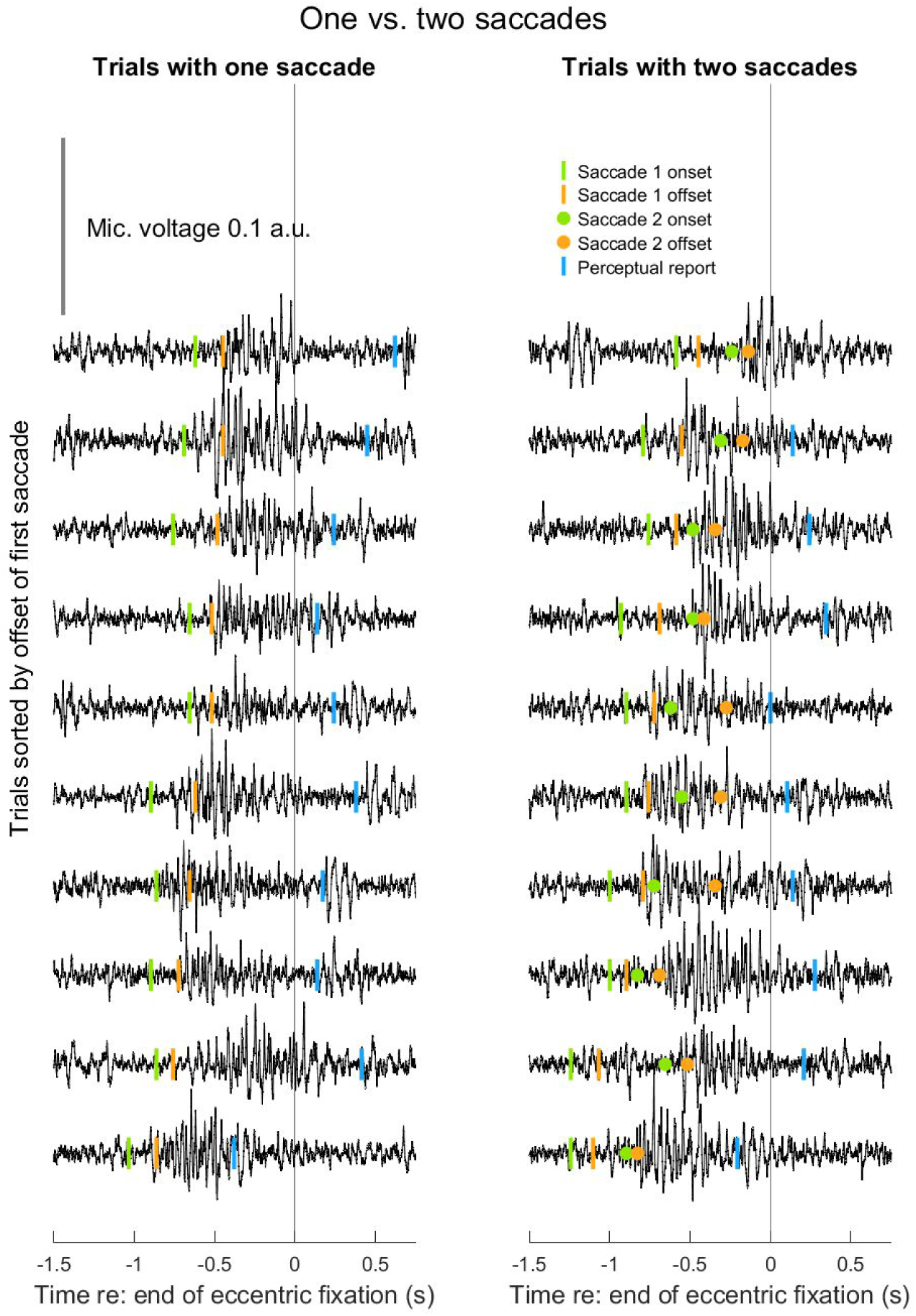
Segmented mic recording corresponding to each reported sound in the self-directed saccade task. The traces are sorted into instances in which there was either one saccade (left) or two saccades (right), and are aligned on the end of eccentric fixation. Vertical sorting is based on the time of offset of the first saccade. The audible sounds last about 1 second and are associated with the eccentric fixation, beginning at the end of the first saccade if only one was made and the end of the second saccade if two were made.

These findings implicate the tensor tympani in the process of linking eye movements and hearing, and illustrate that tensor tympani dysfunction can cause production of *audible* sounds in the ear canal in association with eye movements. But the sounds measured and described above appeared to differ in several ways from the EMREOs we have recorded in participants with normal auditory function. EMREOs are precisely time locked to saccades, and while they extend by about 50-100 ms into periods of steady fixation, they are clearly larger in amplitude during the saccade itself than in the aftermath. However, the temporal imprecision of the video-of-a-video eye tracking method (see Methods) likely precludes even detecting the EMREO itself during the self-directed saccade task described above. To evaluate EMREOs per se, we switched to performance of a visually-guided saccade task with a smaller range of target locations for which we could use quantitative and temporally precise eye tracking and to provide comparison of S98 to our normative dataset of participants with normal auditory function.

### 2. Comparison between EMREOs of S98 and participants without auditory dysfunction

**S98** and N=30 control participants with no known dysfunction of the auditory pathway performed visually-guided saccades in a “grid” task involving a straight ahead fixation and a variety of targets in both horizontal and vertical dimensions (Figure 3). Given that S98 hears audible sounds associated with large leftward eye movements in the left ear, we focused on the EMREOs occurring for leftward saccades and left ears of the normal participants, i.e. pooling across the target locations illustrated in the dashed line box (Figure 3).

**Figure 3.**
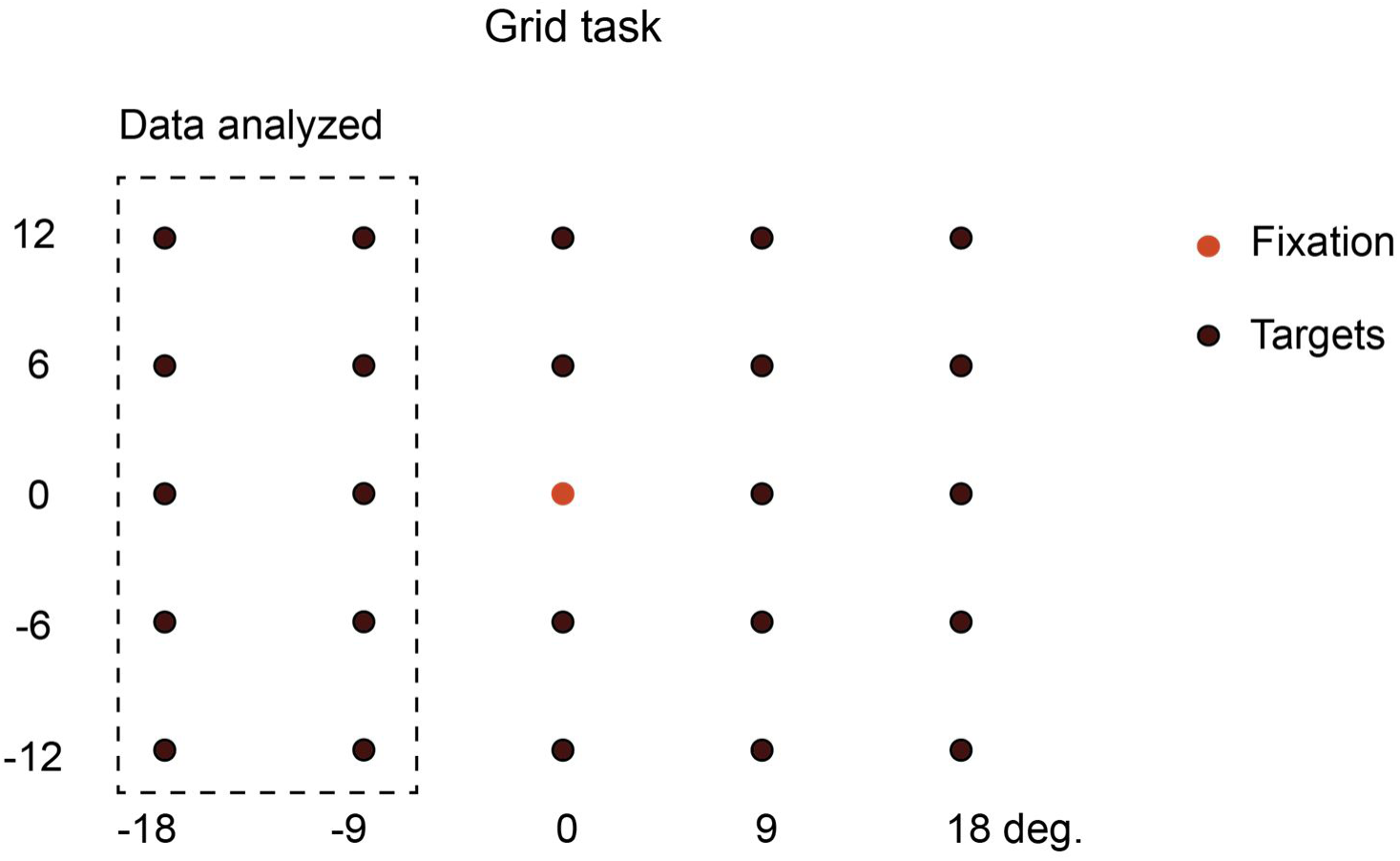
Grid task used for comparison between S98 and N=19 control participants with no known auditory dysfunction. Data were analyzed for the left ears and for leftward saccades, pooling across the target locations contained in the box with the dashed lines.

S98 exhibited a much larger EMREO than population average of the normal participants (Figure 4AB). The wave also began earlier than normal when aligned on saccade onset (Figure 4A). When aligned on saccade offset, the wave was also about 2x larger than in the average of the control group, and large deviations from zero continued for at least 50 ms after saccade offset (Figure 4B). However, the population average of the control group can obscure the results from individual control participants, given that there is individual variation in EMREO timing (King, Lovich et al. 2023). Nevertheless, S98’s peak-trough amplitude was just shy of the 75^th^ percentile (Figure 4C) when this metric was computed for each subject individually.

**Figure 4.**
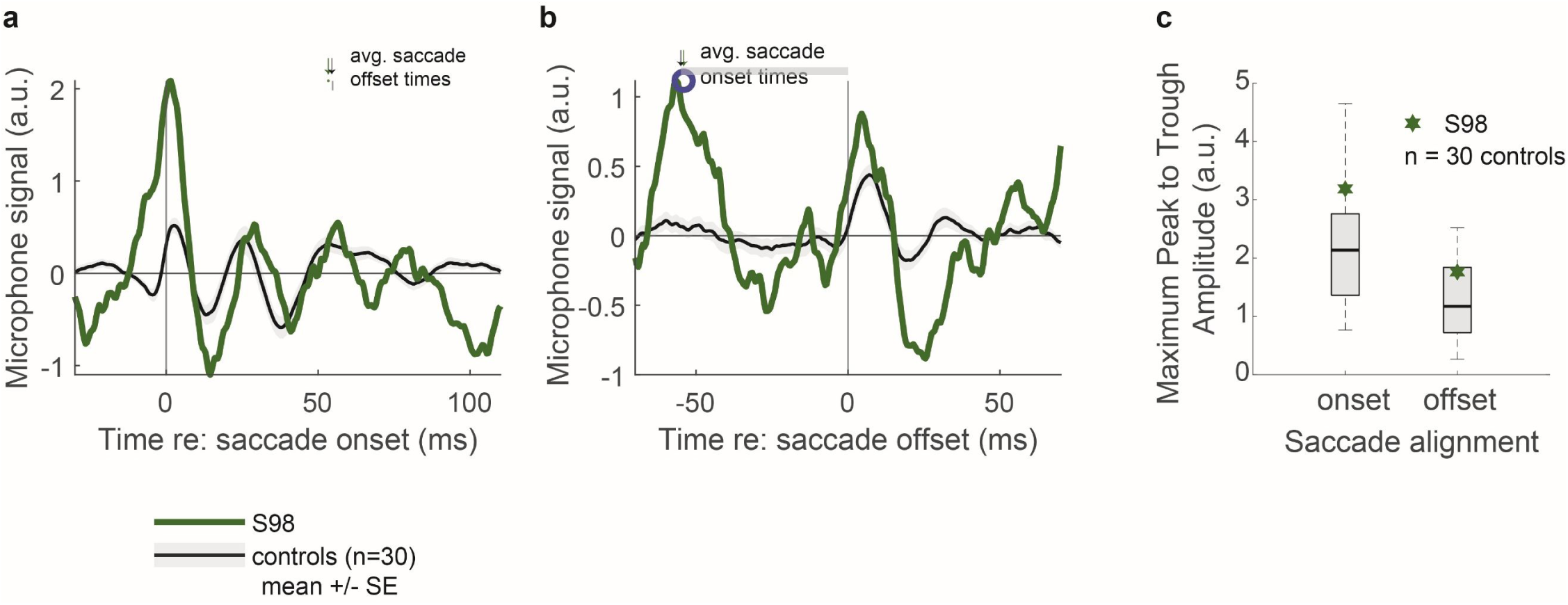
Comparison of S98 EMREOs to a population of typical subjects. Panels a-b show that the EMREO for S98 is dramatically larger than the population mean and SE for both saccade onset (a) and offset-aligned (b) recordings. However, the population average can obscure the range of amplitudes of individual subject waveforms since they can have slightly variable latencies. Panel c controls for this, by depicting (the normal population range of the maximum peak to trough waveform amplitude during the EMREO time window, when aligned to both saccade onset (-5 to 45 ms re: onset) and offset (-5 to 45 ms re: offset). Thick black line = population median, gray box marks the 25^th^-75^th^ percentile range, dashed line identifies the full range of the population response. The max onset-aligned EMREO amplitude for S98 is above the 75^th^ percentile but a few typical subjects have larger EMREO amplitudes. The max offset-aligned EMREO amplitude for S98 is just below the75th percentile.

In short, S98 stood out from the control subjects in having a larger EMREO than is typical, by about a factor of about 50% to 300% depending on the exact metric. However, it remained inaudible to her during performance of this task, suggesting it would need to be larger still to be heard.

## Discussion

In summary, when S98 makes a particularly large saccade to an extreme leftward location, a spasm of the tensor tympani appears to be triggered, producing an audible sound that lasts for nearly a second. The sound requires large eye movements to extreme leftward positions to be elicited. In addition, this participant exhibits classical EMREOs that are larger, occur earlier, and last longer than average, even for saccades in a smaller range that did not produce audible percepts for her.

Since EMREOs were first discovered, considerable attention has focused on both establishing their perceptual purpose as well as their mechanistic underpinnings. Regarding purpose, our theory has been that EMREOs are a signature of the process by which the brain makes the discrepancy between visual and auditory space across eye movements *not* noticeable. Establishing how something produces our ability to perceptually ignore something is a difficult task – most work in this area has looked for evidence of *failures* to compensate for eye movements (e.g. (Lewald and Ehrenstein 1996, Brohl and Kayser 2023, Sotero Silva, Kayser et al. 2025, Sotero Silva, Brohl et al. 2026)), when the big story is that we usually do this quite successfully and don’t notice that our eyes are moving (Boucher, Groh et al. 2001, Metzger, Mullette-Gillman et al. 2004). Indeed, most studies of perceptual impact of eye movements on sound localization or related auditory tasks have reported effects that are small and/or inconsistent in direction – as they should be if our brains are mostly doing a good job at compensating for massive shifts in the orientation of the retina with respect to the head and ears (for review, see (Willett, Groh et al. 2019)).

In the present study, we were able to document a strong association between peripheral auditory mechanisms and eye movements that *did* produce a noticeable perceptual consequence, i.e. a perceived sound when moving the eyes in a certain direction. As noted above, the candidate neural mechanism underlying this percept is a spasm of the tensor tympani muscle, thus connecting one of the potential components creating EMREOs to an overt percept associated with saccades and eccentric fixation.

The involvement of the tensor tympani in EMREOs is the subject of further ongoing work in our group. Unpublished data from our laboratory tentatively indicate that when the tensor tympani is not contributing at all (e.g. when the tendon attaching the tensor tympani to the ossicles has been severed), the EMREO is larger than usual (Lovich, Eliades et al. 2026). Taken together with the present findings, this suggests that a properly functioning tensor tympani normally serves to modulate (and more specifically reduce) the amplitude of the EMREO signal, limiting the likelihood that this signal will be loud enough to be heard. That said, additional unpublished work indicates that the stapedius and outer hair cells also contribute to the generation of this signal (Lovich, Eliades et al. 2026) – the current study should in no way be taken as suggesting that the tensor tympani is the exclusive mechanism by which information about eye movements into peripheral auditory processing, nor do the current findings suggest that mediating eye movement signals need be the only thing that the tensor tympani does.

From the perspective of symptomology, S98 could be considered to have a form of gaze evoked tinnitus. Defined as the percepts of sounds associated with eye movements, gaze-evoked tinnitus has been observed in the context acoustic neuromas and cerebellar pontine angle surgery to remove them (Wall, Rosenberg et al. 1987, Cacace, Lovely et al. 1994, Coad, Lockwood et al. 2001, Lockwood, Wack et al. 2001, Biggs and Ramsden 2002, Albuquerque and Bronstein 2004, Baguley, Phillips et al. 2006), with incidence estimated at 19% in patients undergoing such surgery (Baguley, Phillips et al. 2006). These percepts tend to be heard in the affected ear, and tend to involve chiefly the horizontal dimension, with greater loudness for more eccentric positions toward the affected ear, similar to the pattern observed for S98. However, in at least some cases these percepts tend to have a tonal element, with pitch changing for different eye positions (Cacace, Lovely et al. 1994), which was not observed here. Nevertheless, our study suggests a possible explanation for the experiences of acoustic neuroma/surgical patients: the loss or compromise of auditory input on one side may make it difficult for the brain to segregate signals that are related to eye movements from those that are related to incoming sound, producing a misinterpretation of eye movement and eye position information as actual sound. Indeed, activity related to gaze evoked tinnitus in has been observed in a variety of locations in the brain (Lockwood, Wack et al. 2001, van Gendt, Boyen et al. 2012).

## Methods

All procedures involving human subjects were approved by Duke University’s Campus Institutional Review Board. Methods for recording EMREOs are described in detail in previous publications (King, Lovich et al. 2023, Lovich, King et al. 2023, Lovich, King et al. 2023) and are recapped here.

### Medical background

S98 (age 67 y.o.) reports a diagnosis of tensor tympani myoclonus. Superior semicircular canal dehiscence (SSCD), a common cause for hearing sounds during eye movements, was not considered to be the cause of her complaint, because other than periods of hyperacusis, medical evaluations did not reveal nor did she report any symptoms commonly associated with SSCD (autophony, vestibular problems, hearing loss or fullness in the ear, oscillopsia). Surgical intervention for her tensor tympani myoclonus has not been recommended.

S98 visited the lab on two occasions in the late summer/fall of 2023. We confirmed she had normal hearing (<25 dB HL at 250, 500, 1000, 2000, 4000 and 8000Hz) and normal tympanometry and acoustic reflexes.

### For recordings of the audible eye movement-related myoclonus

the subject was seated in a sound booth facing a large monitor, head positioned in a table-mounted chin rest to limit movement and stabilize head position. Visible targets were placed on the monitor screen to aid in determining the initial fixation and target location that consistently generated the perceptible fluttering sound for the subject. We ascertained that the subject could reliably generate and perceive the sound when making large leftward saccades (initial fixation position [30° H (rightward), 0° V]; target location [-32°, 6°]), resulting in maximum saccade lengths of approximately 62°). The subject reported only hearing the sound during these large leftward eye movements and only in the left ear. Accordingly, the recording microphone (ER10B+) was placed in the left ear and the subject tapped the right ear mic, placed on the eye tracker chinrest table in front of them, every time they heard the fluttering sound.

The eye tracking system (EyeLink 1000) could not quantitatively capture these eye movements between such extreme eccentric positions, but the eye movements were visible on the EyeLink computer monitor nevertheless. Accordingly, we positioned an iPhone camera to video record the eye tracking computer screen. Video and microphone recordings were aligned by recording hand claps at the initiation of the recordings. Based on the combination of the refresh rate on the EyeLink computer (∼70Hz, or a period of 14 ms) and the video frame rate of the iPhone camera (∼29 Hz, period 35 ms), we estimated alignment accuracy to be within ∼50 ms (14 ms + 35 ms = 49 ms).

### Comparison to normal participants

We evaluated this subject’s EMREOs in comparison to participants (n = 30 [20 female, 10 male, 19-54 years, mean = 27.5 years]) with normal hearing and no identified auditory system impairments. All experimental and data analysis procedures were the same for S98 and the control group, except that we only recorded from the left ear in S98 whereas both ears were recorded in the normal participants. Accordingly, comparisons were made with the normal participants’ left ears only. All recordings were conducted in the sound booth as described above. Ear canal recordings were made with the ER10b+. Eye movements were recorded via the EyeLink1000. As described in (King, Lovich et al. 2023, Lovich, King et al. 2023, Lovich, King et al. 2023), saccades were identified on the basis of velocity criterion and trials were discarded if the saccades were excessively curved, too slow, or if there wasn’t a minimum of 200 ms of steady fixation both before and after the saccade.

All subjects performed a grid task as shown in Figure 3 (Lovich, King et al. 2023), with a central visual fixation and saccade targets ranging from +/-18 deg horizontally and +/-12 deg vertically. Given that the audible sounds S98 perceived only occurred in the left ear and only when making large leftward saccades, we focused our analysis on left ear recordings and leftward visual targets for the control subjects as well. Trials involving these targets were pooled to create an average waveform for each subject. It is worth noting that the maximum saccade length for this task was 21.5° and subject S98 did not detect any audible sounds during this task.

## ACKNOWLEDGMENTS

This work was supported by NIH grant DC020363 to JMG and CDK. We thank Stephanie (Schlebusch) Lovich for thoughtful comments.

## Notes

### Competing Interest Statement

The authors have declared no competing interest.

### Summary of Updates

The datapoint for subject 98 in Figure 4c for the saccade offset aligned responses was incorrect due to a data file inclusion error and has been adjusted. The datapoint is now sitting just below the 75th percentile of the control participants. The figure and associated text have been corrected.

## References

Abbasi, H., C. D. King, S. Lovich, B. Roder, J. M. Groh and P. Bruns (2025). “Eye movement-related eardrum oscillations do not require current visual input.” Hear Res 465: 109346.

Albuquerque, W. and A. M. Bronstein (2004). ““Doctor, I can hear my eyes”: report of two cases with different mechanisms.” Journal of neurology, neurosurgery and psychiatry 75(9): 1363–1364.

Baguley, D. M., J. Phillips, R. L. Humphriss, S. Jones, P. R. Axon and D. A. Moffat (2006). “The prevalence and onset of gaze modulation of tinnitus and increased sensitivity to noise after translabyrinthine vestibular schwannoma excision.” Otol Neurotol 27(2): 220–224.

Biggs, N. D. and R. T. Ramsden (2002). “Gaze-evoked tinnitus following acoustic neuroma resection: a de-afferentation plasticity phenomenon?” Clin Otolaryngol Allied Sci 27(5): 338–343.

Boucher, L., J. M. Groh and H. C. Hughes (2001). “Afferent delays and the mislocalization of perisaccadic stimuli.” Vision Research 41(20): 2631–2644.

Brohl, F. and C. Kayser (2023). “Detection of Spatially Localized Sounds Is Robust to Saccades and Concurrent Eye Movement-Related Eardrum Oscillations (EMREOs).” J Neurosci 43(45): 7668–7677.

Bröhl, F. and C. Kayser (2023). “Detection of spatially-localized sounds is robust to saccades and concurrent eye movement-related eardrum oscillations (EMREOs).” Journal of Neuroscience 43(45): 7668–7677.

Bulkin, D. A. and J. M. Groh (2012). “Distribution of eye position information in the monkey inferior colliculus.” Journal of Neurophysiology 107(3): 785–795.

Bulkin, D. A. and J. M. Groh (2012). “Distribution of visual and saccade related information in the monkey inferior colliculus.” Front Neural Circuits 6: 61.

Cacace, A. T., T. J. Lovely, D. F. Winter, S. M. Parnes and D. J. McFarland (1994). “Auditory perceptual and visual-spatial characteristics of gaze-evoked tinnitus.” Audiology 33(5): 291–303.

Caruso, V. C., D. S. Pages, M. A. Sommer and J. M. Groh (2019). “Compensating for a shifting world: A quantitative comparison of the reference frame of visual and auditory signals across three multimodal brain areas.” bioRxiv: 669333.

Caruso, V. C., D. S. Pages, M. A. Sommer and J. M. Groh (2021). “Compensating for a shifting world: evolving reference frames of visual and auditory signals across three multimodal brain areas.” J Neurophysiol 126(1): 82–94.

Coad, M. L., A. Lockwood, R. Salvi and R. Burkard (2001). “Characteristics of patients with gaze-evoked tinnitus.” Otol Neurotol 22(5): 650–654.

Cohen, Y. E. and R. A. Andersen (2000). “Reaches to sounds encoded in an eye-centered reference frame.” Neuron 27(3): 647–652.

Fu, K. M., A. S. Shah, M. N. O’Connell, T. McGinnis, H. Eckholdt, P. Lakatos, J. Smiley and C. E. Schroeder (2004). “Timing and laminar profile of eye-position effects on auditory responses in primate auditory cortex.” J Neurophysiol 92(6): 3522–3531.

Groh, J. M., A. S. Trause, A. M. Underhill, K. R. Clark and S. Inati (2001). “Eye position influences auditory responses in primate inferior colliculus.” Neuron 29(2): 509–518.

Gruters, K. G., D. L. K. Murphy, C. D. Jenson, D. W. Smith, C. A. Shera and J. M. Groh (2018). “The eardrums move when the eyes move: A multisensory effect on the mechanics of hearing.” Proc Natl Acad Sci U S A 115 (6): E1309–E1318.

Hartline, P. H., R. L. Vimal, A. J. King, D. D. Kurylo and D. P. Northmore (1995). “Effects of eye position on auditory localization and neural representation of space in superior colliculus of cats.” Exp Brain Res 104(3): 402–408.

Hong, F., S. Badde and M. S. Landy (2021). “Causal inference regulates audiovisual spatial recalibration via its influence on audiovisual perception.” PLoS Comput Biol 17(11): e1008877.

Jay, M. F. and D. L. Sparks (1984). “Auditory receptive fields in primate superior colliculus shift with changes in eye position.” Nature 309(5966): 345–347.

Jay, M. F. and D. L. Sparks (1987). “Sensorimotor integration in the primate superior colliculus. I. Motor convergence.” J Neurophysiol 57(1): 22–34.

Jay, M. F. and D. L. Sparks (1987). “Sensorimotor integration in the primate superior colliculus. II. Coordinates of auditory signals.” J Neurophysiol 57(1): 35–55.

King, C. D. and J. M. Groh (2026). “Saccade-related sound pulses and phase resetting contribute to eye movement-related eardrum oscillations (EMREOs).” BioRxiv 2026.03.25.714060.

King, C. D., S. N. Lovich, D. L. Murphy, R. Landrum, D. Kaylie, C. A. Shera and J. M. Groh (2023). “Individual similarities and differences in eye-movement-related eardrum oscillations (EMREOs).” Hear Res 440: 108899.

Kording, K. P., U. Beierholm, W. J. Ma, S. Quartz, J. B. Tenenbaum and L. Shams (2007). “Causal inference in multisensory perception.” PLoS One 2(9): e943.

Lee, J. and J. M. Groh (2012). “Auditory signals evolve from hybrid-to eye-centered coordinates in the primate superior colliculus.” Journal of Neurophysiology 108(1): 227–242.

Leon, M. E., C. Ramos and P. E. Maldonado (2026). “Eye Movement-Related Eardrum Oscillations (EMREOs) Occur without Visual Input But Are Reduced during Closed Eyelids.” J Neurosci 46(2).

Lewald, J. and W. H. Ehrenstein (1996). “The effect of eye position on auditory lateralization.” Exp Brain Res 108(3): 473–485.

Lockwood, A. H., D. S. Wack, R. F. Burkard, M. L. Coad, S. A. Reyes, S. A. Arnold and R. J. Salvi (2001). “The functional anatomy of gaze-evoked tinnitus and sustained lateral gaze.” Neurology 56(4): 472–480.

Lovich, S., S. Eliades, D. Kaylie, C. King, C. Shera and J. M. Groh (2026). “ Mechanisms for generation and control of EMREOs (Eye movement-related eardrum oscillatons)..” Association for Research in Otolaryngology Midwinter Meeting, San Juan PR.

Lovich, S. N., C. D. King, D. L. Murphy, R. Landrum, C. A. Shera and J. M. Groh (2023). “Parametric information about eye movements is sent to the ears.” Proceedings of the national academy of sciences 120(48): p. e2303562120.

Lovich, S. N., C. D. King, D. L. K. Murphy, H. Abbasi, P. Bruns, C. A. Shera and J. M. Groh (2023). “Conserved features of eye movement related eardrum oscillations (EMREOs) across humans and monkeys.” Philos Trans R Soc Lond B Biol Sci 378(1886): 20220340.

Maier, J. X. and J. M. Groh (2010). “Comparison of gain-like properties of eye position signals in inferior colliculus versus auditory cortex of primates.” Frontiers in Integrative Neuroscience 4: 121–132.

Metzger, R. R., O. A. Mullette-Gillman, A. M. Underhill, Y. E. Cohen and J. M. Groh (2004). “Auditory saccades from different eye positions in the monkey: implications for coordinate transformations.” Journal of Neurophysiology 92(4): 2622–2627.

Mullette-Gillman, O. A., Y. E. Cohen and J. M. Groh (2005). “Eye-centered, head-centered, and complex coding of visual and auditory targets in the intraparietal sulcus.” Journal of Neurophysiology 94(4): 2331–2352.

Mullette-Gillman, O. A., Y. E. Cohen and J. M. Groh (2009). “Motor-related signals in the intraparietal cortex encode locations in a hybrid, rather than eye-centered, reference frame.” Cerebral Cortex 19(8): 1761–1775.

Populin, L. C., D. J. Tollin and T. C. Yin (2004). “Effect of eye position on saccades and neuronal responses to acoustic stimuli in the superior colliculus of the behaving cat.” J Neurophysiol 92(4): 2151–2167.

Porter, K. K., R. R. Metzger and J. M. Groh (2006). “Representation of eye position in primate inferior colliculus.” Journal of Neurophysiology 95(3): 1826–1842.

Rohe, T. and U. Noppeney (2015). “Cortical hierarchies perform Bayesian causal inference in multisensory perception.” PLoS Biol 13(2): e1002073.

Russo, G. S. and C. J. Bruce (1994). “Frontal eye field activity preceding aurally guided saccades.” Journal of Neurophysiology 71(3): 1250–1253.

Sotero Silva, N., F. Brohl and C. Kayser (2026). “Sound lateralization ability is affected by saccade direction but not eye movement-related eardrum oscillations.” J Neurophysiol 135(4): 1085–1097.

Sotero Silva, N., C. Kayser and F. Brohl (2025). “Unraveling eye movement-related eardrum oscillations (EMREOs): how saccade direction and tympanometric measurements relate to their amplitude and time course.” Hear Res 461: 109276.

Stricanne, B., R. A. Andersen and P. Mazzoni (1996). “Eye-centered, head-centered, and intermediate coding of remembered sound locations in area LIP.” Journal of Neurophysiology 76(3): 2071–2076.

van Gendt, M. J., K. Boyen, E. de Kleine, D. R. Langers and P. van Dijk (2012). “The relation between perception and brain activity in gaze-evoked tinnitus.” J Neurosci 32(49): 17528–17539.

Wall, M., M. Rosenberg and D. Richardson (1987). “Gaze-evoked tinnitus.” Neurology 37(6): 1034–1036.

Werner-Reiss, U., K. A. Kelly, A. S. Trause, A. M. Underhill and J. M. Groh (2003). “Eye position affects activity in primary auditory cortex of primates.” Current Biology 13: 554–562.

Willett, S. M., J. M. Groh and R. K. Maddox (2019). Hearing in a “moving” visual world: Coordinate transformations along the auditory pathway. Springer Handbook of Auditory Research: Multisensory Processes The Auditory Perspective. A. K. C. Lee, M. T. Wallace, A. B. Coffin, A. N. Popper and R. R. Fay, Springer. 68.

Zella, J. C., J. F. Brugge and J. W. Schnupp (2001). “Passive eye displacement alters auditory spatial receptive fields of cat superior colliculus neurons.” Nat Neurosci 4(12): 1167–1169.

Zwiers, M. P., H. Versnel and A. J. Van Opstal (2004). “Involvement of monkey inferior colliculus in spatial hearing.” J Neurosci 24(17): 4145–4156.

